# A Parallelized, Automated System for Higher-Throughput Spatial Navigation Testing of Rats

**DOI:** 10.1101/2025.07.22.666070

**Authors:** Violet T. Saathoff, Josiah A. Quinn, Zhana D. Prince, Charles I. O’Leary, Anna K. Gillespie

## Abstract

Spatial navigation tasks for rodents have been an extremely useful tool for studying cognitive processes for decades, but are often laborious to administer and sensitive to experimenter-induced variability. To address these limitations, we have developed a parallelized maze system which runs four highly flexible maze environments simultaneously with automated reward delivery, enabling reduced experimenter involvement, increased throughput, and streamlined data collection. Further, our software tracks subject performance and manages the progression through a pre-set curriculum as subjects learn, reducing experimenter error and ensuring comparable learning curves across cohorts. Equipped with IR beam-break based reward ports, peristaltic pumps for liquid reward delivery, and overhead video tracking, our current maze accessories can be flexibly reconfigured to enable a vast array of behavioral tasks and support a wide range of experimental questions. Pilot cohorts of young (4-8 month) and aged (28-32 month) male and female rats demonstrate that this apparatus enables the efficient training of a series of increasingly difficult visually cued and memory-dependent spatial navigation task variants across a wide range of performance abilities.

**Significance Statement:** To increase the efficiency and reliability of spatial navigation task administration, we have designed a behavioral apparatus which enables the training of four subjects in parallel on a variety of tasks with automated reward delivery. Further, we developed software to track subject performance and advance training parameters through a pre-set curriculum to reduce experimenter error and enable robust comparison of learning metrics across cohorts.

## 1. Introduction

For decades, scientists have studied the behavior of rodents navigating maze-like environments in order to understand diverse cognitive processes such as memory, decision-making, inference, fear, regret, sociability, and many more (Morris 1981; Olton and Samuelson 1976; Steiner and Redish 2014; Tolman 1948; Wijnen et al. 2024). Spatial navigation tasks for rodents have been particularly powerful tools due to their ethological relevance — a hungry rat will reliably seek food in a complex environment, just as it would in the wild. Spatial navigation tasks can provide a rich and sensitive behavioral readout of neural computations because they often incorporate multiple cognitive components: for instance, a subject might need to identify its location in an environment, recall past locations of food, obstacles, or threats, identify the most valuable location, plan a route towards that goal, execute the movement, and evaluate the outcome of that choice (Wijnen et al. 2024). Depending on the task, this might require using multiple types of memory — declarative, procedural, and/or working memory. Due to their sensitivity and reliability, spatial navigation tasks have become the gold standard for characterizing cognitive impairments in the case of neurodegenerative and other disease models (Astur et al. 2002; D’Hooge and De Deyn 2001; Vorhees and Williams 2014). Clever experimental design can allow researchers to tease apart the different cognitive components of spatial navigation challenges, and by incorporating simultaneous neural recording approaches, researchers have been able to begin describing the neural mechanisms that underlie these cognitive processes. Perhaps the most well known finding stemming from this approach was the discovery of spatially tuned neurons in the hippocampus (O’Keefe and Dostrovsky 1971), which has since inspired decades of research into the neural activity underlying the encoding of spatial experiences and their subsequent storage as memory (Findlay et al. 2020; Moser et al. 2015). However, there is still much that we don’t understand about how the brain enables spatial navigation behavior. Thus, many further experiments, involving creative task variants, different neural recording modalities, and cohorts of different ages, sexes, and disease conditions remain a high priority.

However, spatial navigation tasks have some notable limitations which can slow progress. They are typically large and often designed for a single behavioral task, limiting the utility of each piece of equipment. Subjects are usually run serially, limiting behavioral cohort sizes and making behavioral testing slow and laborious. A human experimenter is often an essential part of the training/experimentation process (for example, delivering a food reward or providing an escape opportunity in an aversive condition), adding an experimental variable which is hard to track and which limits the scalability of behavioral testing (Chesler et al. 2002; Pioli et al. 2014). When combined with neurophysiology, the need for additional subject space, tethering, or overhead tracking can place additional constraints on maze environments. Finally, because these experiments can be so costly and time-consuming, subjects are often selected for their motivation and aptitude for the task to maximize the chance of successful data collection. As a result, these cohorts may be biased towards high performers rather than reflecting the extent of natural variability in the population.

We set out to develop a spatial navigation maze apparatus that would address many of these limitations. Specifically, we set out to meet the following five criteria. First, we wanted to be able to run multiple subjects at once in order to increase efficiency, enabling larger cohort sizes (Hao et al. 2021; Poddar et al. 2013). Second, we wanted to automate the task in order to minimize experimenter-induced variability (Chesler et al. 2002; Pioli et al. 2014; Qiao et al. 2019; Zou and Li 2019). Third, we wanted the apparatus to be flexible enough to run a variety of tasks (Porter et al. 2025). Not only would this provide more opportunities to use the same equipment for a range of experimental questions, but it would allow us to design a task curriculum that spans a range of difficulty, progressing from an “easy” task variant to more challenging ones so that we could run structured, systematic behavioral testing across the full range of individual abilities (Holleman et al. 2019). Fourth, our apparatus needed to be compatible with wireless neurophysiological recordings. For hippocampal recordings to be analyzed in relation to spatial location, this required overhead position tracking as well as sufficiently wide corridors for an implanted rat. The training curriculum (including initial task acquisition) would ideally take less than 4 weeks, a time window similar to our periods of high quality neural recording signal. Finally, our behavioral task also needed to be highly structured, to enable the collection of many comparable behavioral trials and ensure good spatial coverage of the entire environment for subsequent analysis.

Our resultant maze apparatus is a low-cost, scalable, highly automated, and parallelized setup that can be configured to run a variety of different spatial navigation tasks. Our proof-of-principle apparatus allows experimenters to administer spaced behavioral training sessions to four rodent subjects simultaneously. The behavioral arenas described here are designed to be consistent, efficient, and simple to construct. The logic of the task can be easily customized; here we focus on a task which includes both visually cued and memory-dependent variants. We have developed a python package that automates the advancement through a series of task variants for each subject based on set performance criteria. This highly structured, automated training program reduces the opportunities for experimenter error and ensures that all subjects attain certain criteria before advancing, providing comparable behavioral epochs within which to study neural mechanisms of learning and performance. As much as possible, our system relies on open-source software packages and hardware designs. All custom designs, components, and software are shared here as a resource (Appendix B), in addition to extensive instructions and part lists, to make this system easy to scale, replicate, and implement.

## 2. Materials and Methods

### 2.1. Design Overview

The maze apparatus consists of four fully enclosed 6-arm maze environments overlooked by a centralized control station (Figure 1A,B, Figure 1-1A). Each maze environment consists of a central home area with a single reward port, and 6 radial arms each with a reward port at each arm end (Figure 1C). Each maze is equipped with an overhead camera, a Pumps and Electronics Rack (PER), and a drainage system to handle liquid waste. At the control station, the experimenter can view the overhead video from each maze, control the behavior of the task, and monitor performance in real time (Figure 1-1 B). From inside the maze, subjects can view both local intramaze cues as well as distal cues on the ceiling, room walls, and floor due to the transparent acrylic floor, ceiling, and mesh arm lids (Figure 1-1C). Whether the distal cues are used to guide behavior is outside the scope of this study, but could easily be tested in future by limiting the external view through the walls and ceiling with contact paper or other means of visual obstruction.

**Figure 1:**
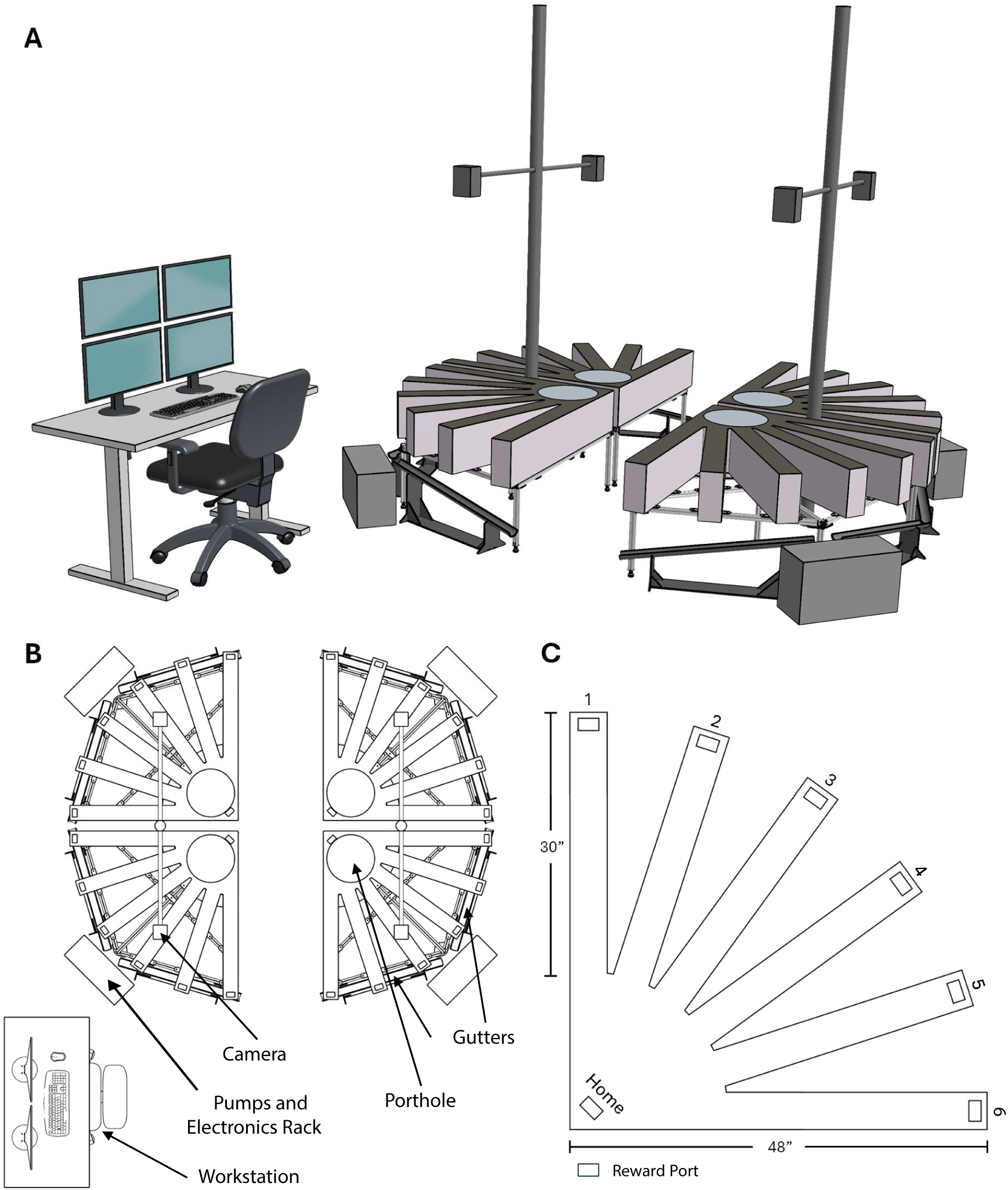
Room Overview: A 3D view of the entire room. Note that colors/opacity of various components are not true to the actual design, and are instead intended to help with visual clarity. B Top-down view of the entire room, showing the layout of all four maze environments, as well as the workstation (lower left). C Top-down schematic view of the maze environment, with reward ports indicated by rectangles.

**Figure 1-1:**
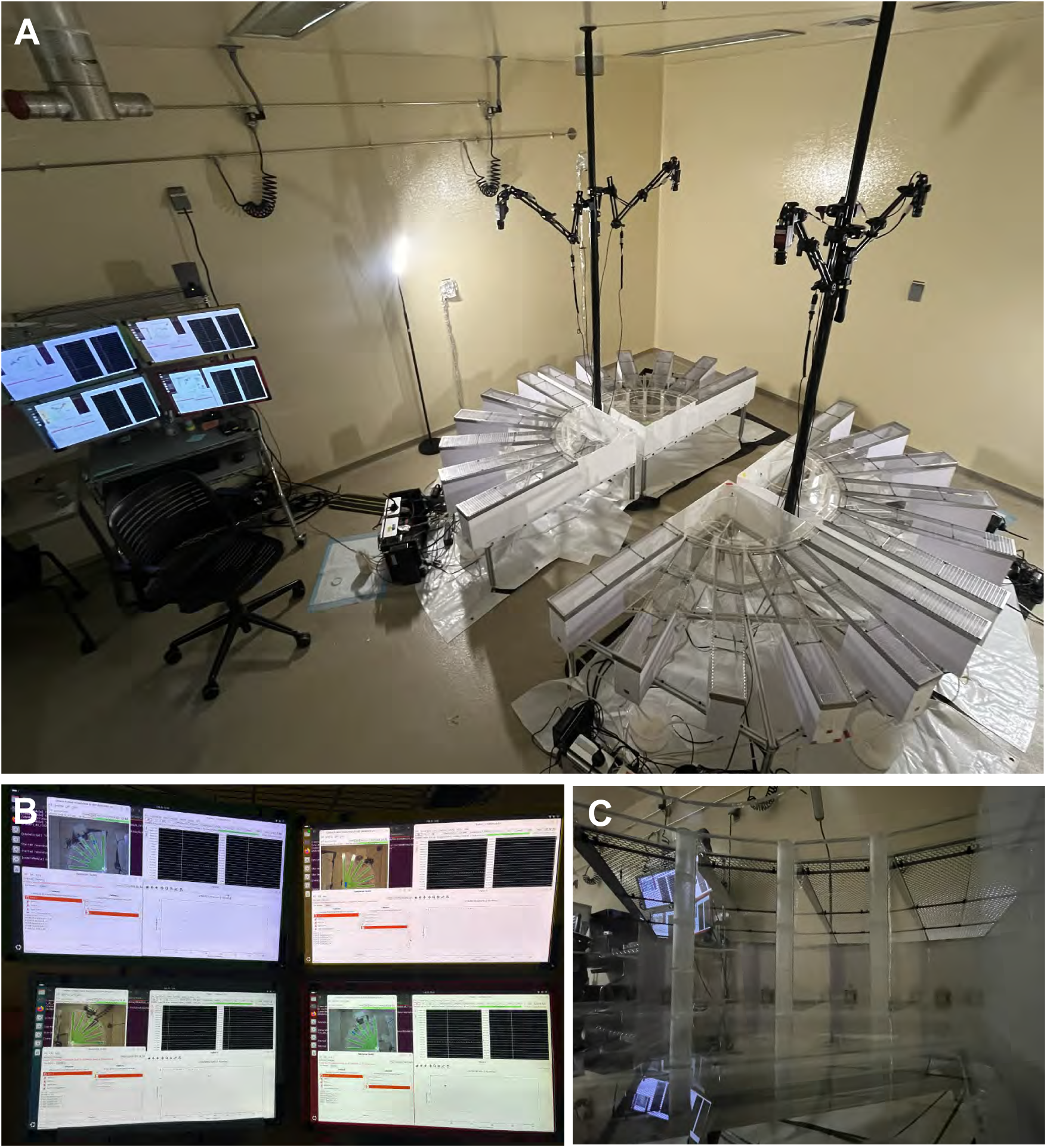
Room Overview: A Photo of the entire room. B Experimenter view of the 4 monitors, one per maze. On each monitor, the experimenter can view the overhead camera video, a live plot of behavioral performance, streaming behavioral variables in the SpikeGadgets Trodes software, and the SpikeGadgets StateScript behavioral control module. C Rat’s-eye view from inside the maze closest to the command station, looking outward from the home port towards the desk/closest wall. Note the clear view through the maze lids of distal cues (the ceiling and walls) as well as features within the maze (8020 below the clear floor, corners, and ports at the end of each arm).

### 2.2. Maze Construction

The environment is primarily constructed out of laser cut acrylic (Figure 2A,B) which rests on an 8020 extruded aluminum base (Figure 2C). Acrylic was selected because it comes in a variety of colors (both transparent and opaque), and because we can manufacture and assemble the parts in house easily and consistently. Where pieces are joined at 90*^◦^* angles, box joints were used to carefully and reliably control the positions of various components. The floor of the maze is made out of 3/16” clear acrylic which provides satisfactory rigidity. The transparent floor did not appear to cause any anxiety in rat subjects; they freely explore immediately when placed in the environment and did not show any preference for maze areas with an opaque floor. Maze walls were made from 1/8” acrylic, which provides sufficient rigidity. Most of the walls were made from clear acrylic to create a complex visual environment, although adding matte contact paper to around the ends of each arm was helpful for minimizing potentially confusing reflections of illuminated reward ports. The outer walls were made from white acrylic to prevent the subjects from seeing each other.

**Figure 2:**
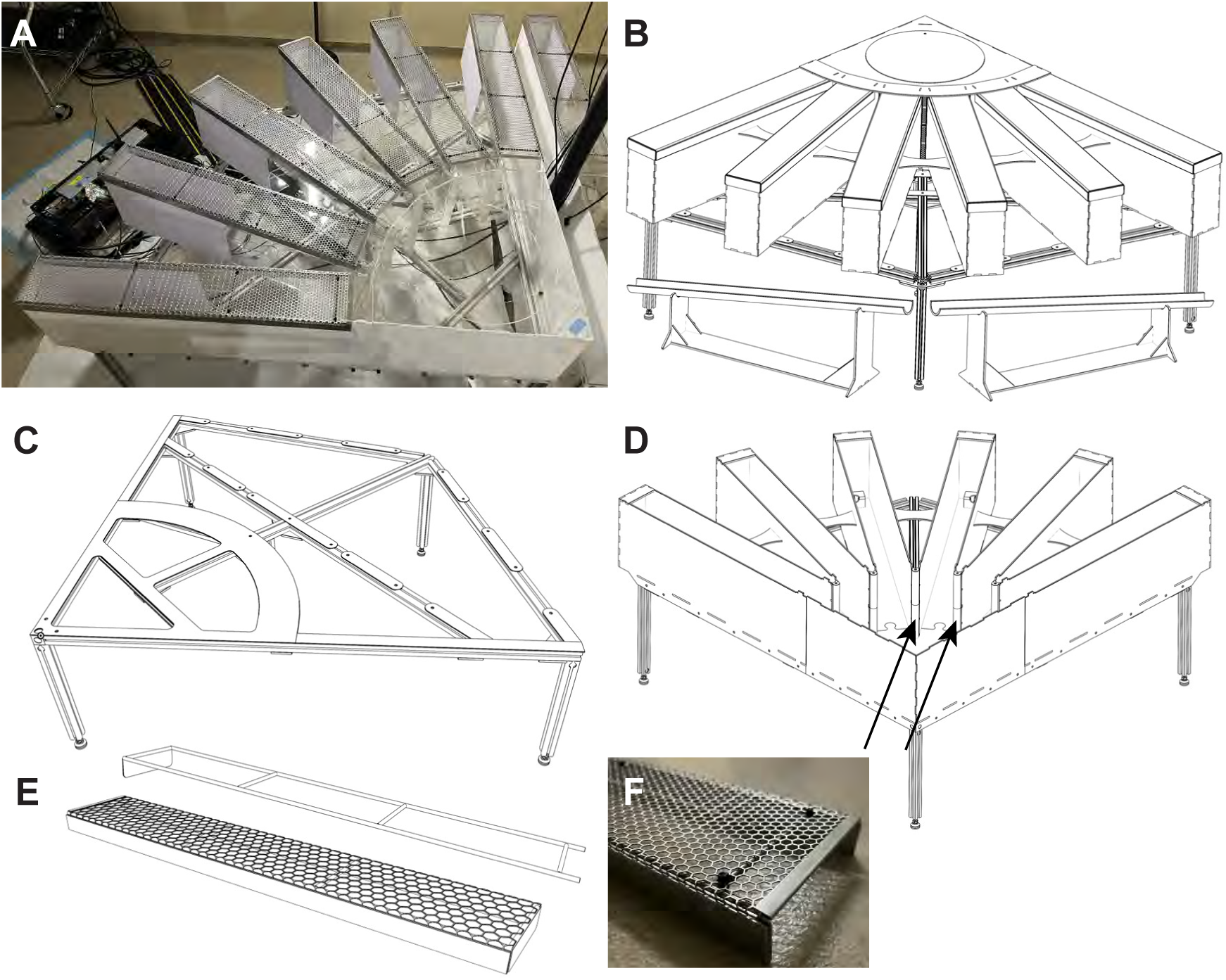
6-Arm Maze: A Photograph of the fully-assembled maze (side) with lids and ceiling included. B 3D view of the fully-assembled maze (back), with the drainage gutters clearly visible. C 3D view of the base (side). The base is made from 8020 extruded aluminum, and has laser-cut, 1/16” acrylic tabs bolted to it. These tabs are then glued to the floor of the maze to help secure it to the 8020 base. D 3D view of the maze (front) with the lids and ceiling removed. Note the 3D-printed corner pieces (arrows) which join the walls between adjacent arms. E 3D view of the frame for an arm lid (top) and the complete arm lid (bottom). The frame adds rigidity and robustness to the hexagonal metal mesh lids, which helps to prevent them from becoming damaged over time due to normal use. Besides this panel, lids are displayed as being solid when in reality they are made of a hexagonally-perforated aluminum affixed to a stainless steel frame. F Photograph of the maze lids to show the texture of the hexagonal mesh.

The environment is fully enclosed to prevent subjects from escaping, with a ceiling height of 9”. The ceiling over the home area is made of clear 3/16” acrylic, with a 10.5” porthole for putting subjects into the maze and a clear 1/8” acrylic porthole cover (Figure 2A,B). The ceiling over each arm is a removable hexagonally perforated aluminum lid, affixed onto a stainless steel rod frame (Figure 2A,E,F). The perforated lids allow for air to enter the environment, introduce only minimal occlusion for overhead camera tracking, and are easily removable to allow the experimenter to access maze arms when needed; all exposed perforations were hemmed to protect both the experimenter and the subject from sharp edges. To secure the lids, one lid end slides under an overhang in the ceiling of the home area, while the outer end has an end cap added. While a fully-enclosed design does prevent tethered recordings, minimizing experimenter supervision was critical for successfully running parallelized behavior.

For the maze base, we selected 8020 extruded aluminum framing as it is modular, strong, and easily attainable; easily supporting the maze while giving us options for routing tubing, cables, and drainage (Figure 2C). To prevent undue stresses in the acrylic floor which could cause eventual cracking, care was taken to ensure the outer rails of the frame were positioned as close to 90*^◦^* as possible, and that the feet of the 8020 base distributed the weight of the maze as evenly as possible. To secure the maze to the base, the main body of the maze was epoxied to a layer of 1/16” acrylic, which was then bolted to the 8020 base (Figure 2C), additionally, the outer walls of the maze were themselves bolted directly to the 8020 base, providing a very solid connection between the base and the maze. Acrylic pieces were bonded using a clear, slow-cure 2-part epoxy (BSI-205), or occasionally with acrylic cement (Weld-On 3). For better gluing surfaces, the walls of adjacent arms are joined with 3D-printed corner pieces (Figure 2D, Figure 1-1C).

### 2.3. Reward Ports

The most critical features of the maze are the reward ports, which dispense a liquid food reward (evaporated milk with 5% sucrose added). Each port consists of an IR photogate (emitter and receiver; 890nm), a white flatmount LED, an 1/8” inner diameter inlet port, a 1/4” inner diameter outlet port, and an RJ-45 port for communication with the ECU (Figure 3A-D). The reward ports’ body and their electronics are based on an established design (Roumis 2020). Compared to the original design, barbed tube fittings were added to make it easier to connect tubing to the reward ports, the drainage port was enlarged to help avoid overflows during cleaning, and the footprint was streamlined to remove crevices where refuse could easily accumulate. The body of the reward ports were printed on a FormLabs Form 3 printer using clear resin and a custom PCB is populated and installed inside. Printing with clear resin allows the white LED to shine through the translucent body of the reward port, allowing the port to illuminate and serve as an easily visible cue. When a subject breaks the IR beam, a signal is recorded and can trigger the delivery of liquid through the port via an associated peristaltic pump.

**Figure 3:**
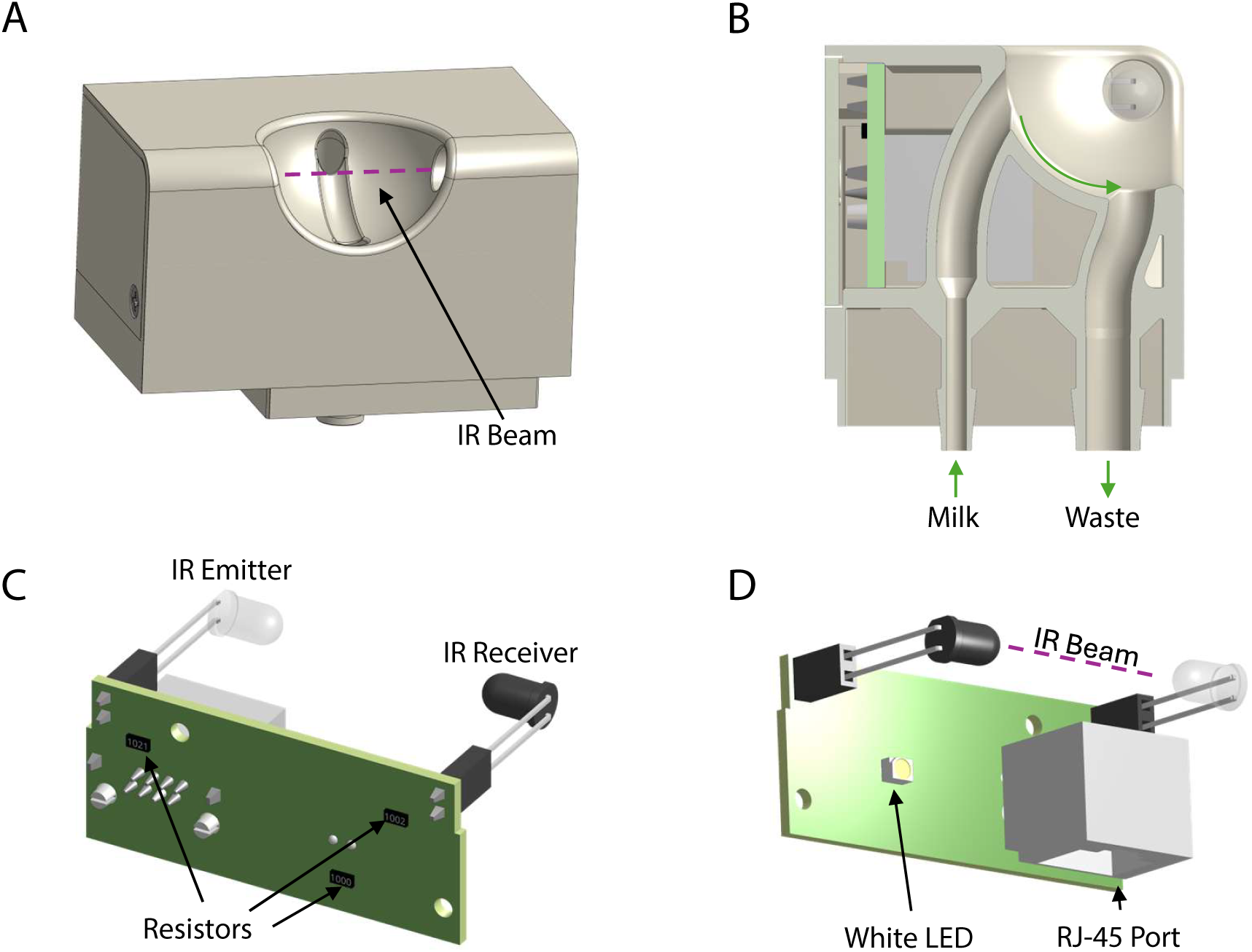
Reward Port: A Reward port adapted from (Roumis 2020), front view. When the rat’s nose enters the cavity, the IR beam is broken, and the port is “triggered”. This behavioral event can trigger milk delivery and/or other environmental actions. B Cutaway view, showing the flow of liquid through the reward port. Milk enters the port through an 1/8” tube, and drains out through a 1/4” tube, both connected to the well with a barb fitting. C Reward port PCB, back view. D Reward port PCB, 3D front view. The RJ-45 port allows the port to be connected to the pump control electronics via an ethernet cable, enabling readout of the IR beam state and control over the state of the white LED.

### 2.4. Pumps and Electronics Rack (PER)

Each maze includes several key electronic accessories, which we combine into a single organized 4U server rack and refer to as the pumps and electronics rack (PER; Figure 4A). The PER houses a compact computer (Intel NUC), a router for connecting to the overhead cameras, a pump control/reward port breakout board, and peristaltic pumps for liquid reward delivery. It also houses a neural data acquisition system and environmental control unit. We chose products from SpikeGadgets (an ECU and either a Main Control Unit or LoggerDock) for their price point and data collection capabilities, but other systems such as OpenEphys (Siegle et al. 2017), Arduino, BControl (*Bcontrol* 2025), or Bonsai (Lopes and Monteiro 2021) environmental control strategies could be used. Each PER is connected to the central command station via the NUC, which sends display and peripheral signals to a KVM switch so that a single keyboard/mouse can control any of the maze stations. The PER for each maze is located close to the maze to minimize tubing/cable lengths, and to minimize the tripping hazard that tubing and cables can cause. A custom mounting plate for the pumps and the pump control PCB was designed to mount to the server rack and secure all pumps, mitigating any vibration caused by pump activity.

**Figure 4:**
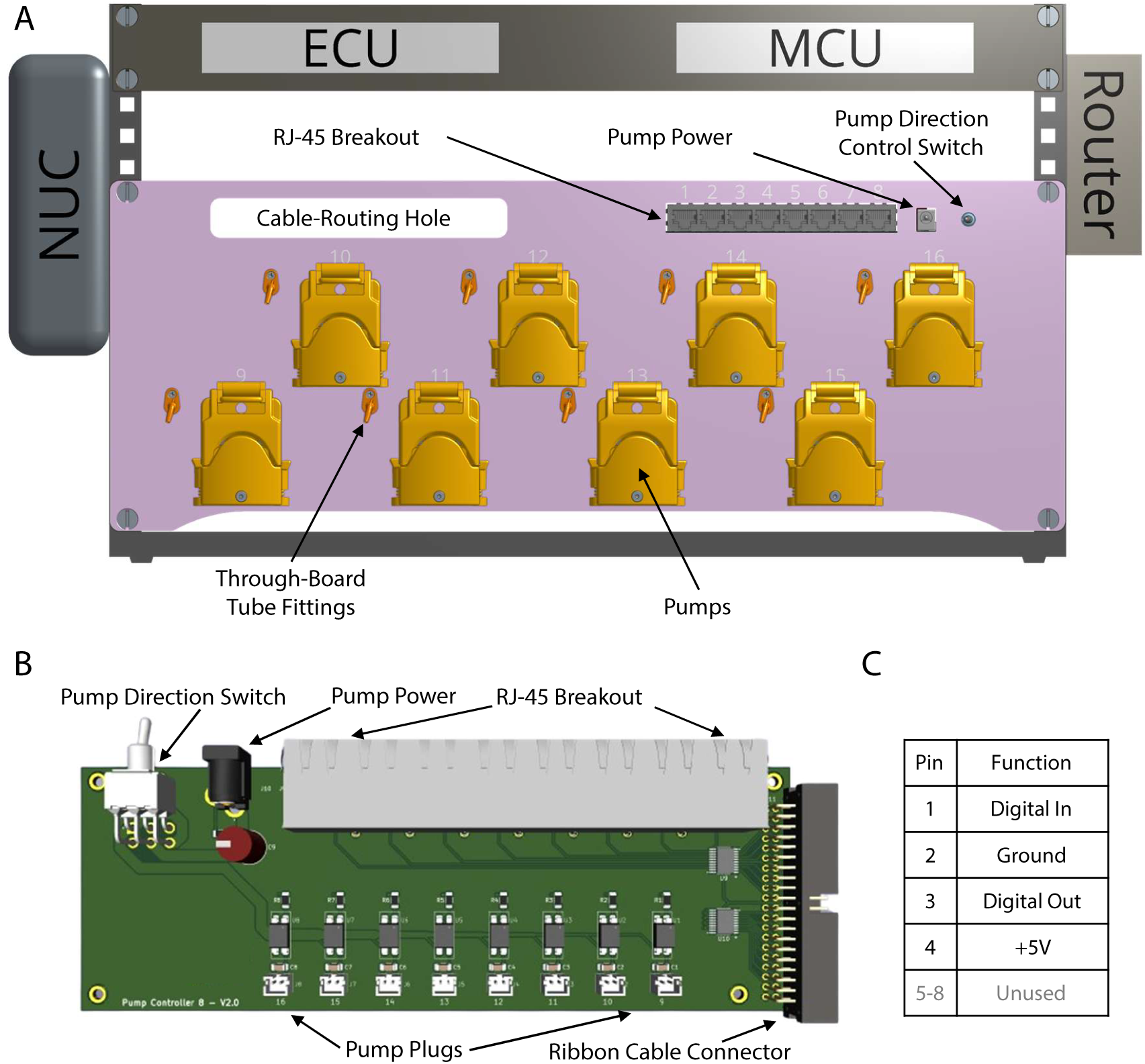
Pumps and Electronics Rack (PER): A Front-view of the Pumps and Electronics Rack (PER); a server rack which integrates the computer (NUC), the SpikeGadgets Main Control Unit (MCU), the SpikeGadgets Environmental Control Unit (ECU), and Pump Plate (pink). The Pump Plate, made from 4mm anodized aluminum, can hold up to 8 Boxer 9QX-9007.730 peristaltic pumps, and mounts a Pump Controller PCB. The pump control electronics also include an RJ-45 breakout board which routes reward port and camera sync signals. Tubing (not shown) routes liquid from a single 1L reservoir to each of the pumps, then through the pump plate via the through-board tube fittings to the reward ports. B Top-down view of the pump controller/RJ-45 breakout PCB. Each board connects to the ECU via a ribbon cable, and can individually power up to 8 pumps via the JST 2-PH connectors. Each board can also connect to 8 accessories (typically reward ports) via ethernet cables. C Pinout for each RJ-45 breakout port. When a breakout port is connected to a reward port, pin 1 represents the state of the IR beam, pin 3 controls the output of the white LED, and pin 4 powers both the IR LED, and the IR receiver.

**Figure 4-1:**
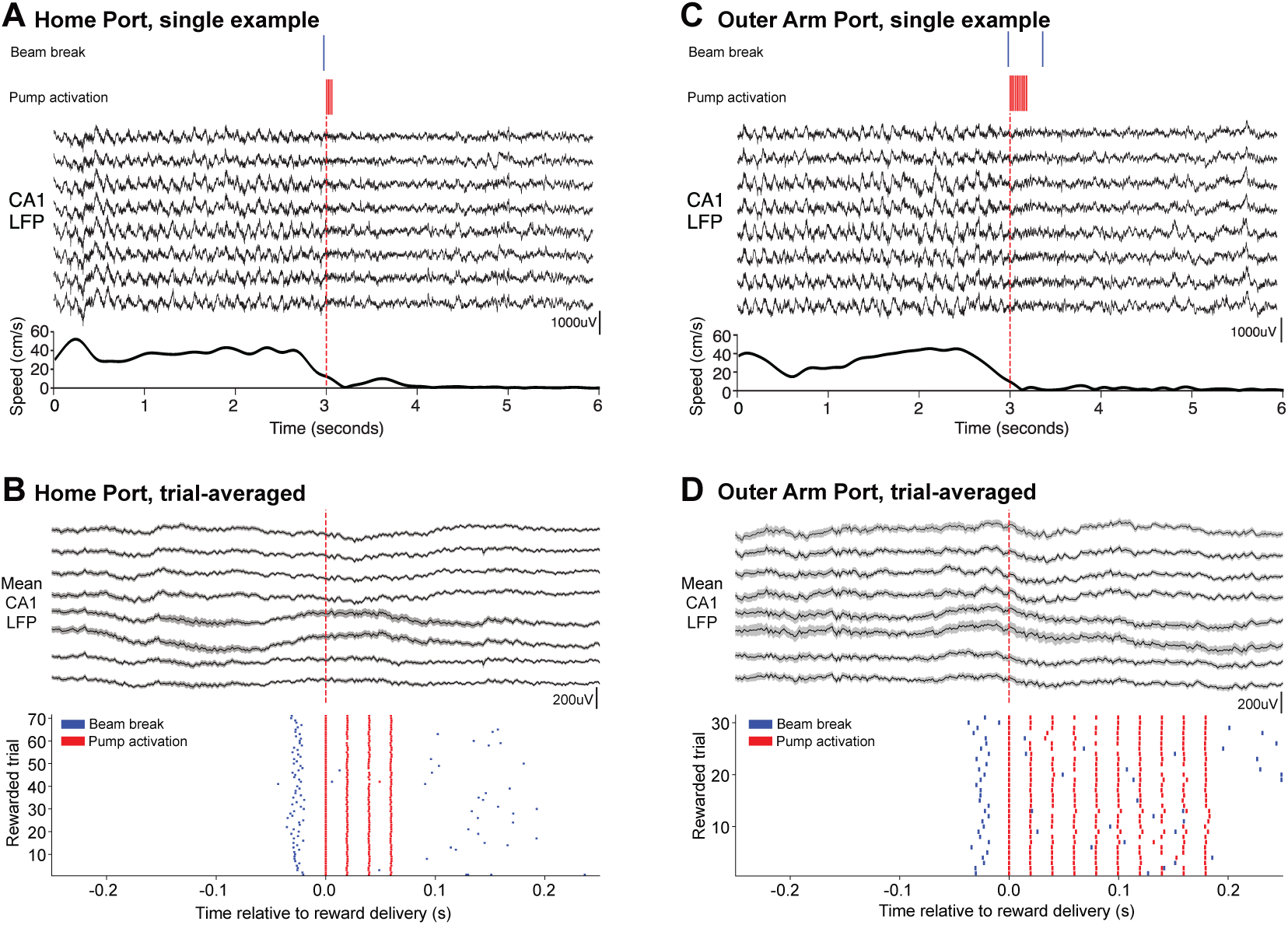
Electrophysiology During Maze Behavior: A Top: single example of hippocampal LFP traces (tetrodes targeted to dorsal CA1 cell layer) during maze behavior, aligned to the onset of reward delivery at the home port. Bottom: rat speed over this time period, showing deceleration prior to and stopping at the reward port. B Top, hippocampal LFP traces averaged over all home port visits within a behavioral epoch, aligned to the onset of reward delivery. Bottom, raster plot illustrating the time of beam break (blue) and reward delivery (red) across all home port visits. C Top: single example of hippocampal LFP traces (tetrodes targeted to dorsal CA1 cell layer) during maze behavior, aligned to the onset of reward delivery at the outer arm ports. Bottom: rat speed over this time period, showing the deceleration and stopping at the reward port. D Top, hippocampal LFP traces averaged over all outer arm port visits within a behavioral epoch, aligned to the onset of reward delivery. Bottom, raster plot illustrating the time of beam break (blue) and reward delivery (red) across all outer arm port visits. Data visualization enabled by Figpack (Magland 2025)

We use peristaltic pumps to deliver small amounts of liquid reward to reward ports. After extensive testing of many pump models, we selected Boxer 9QX peristaltic pumps. When delivering rewards, the ECU sends a TTL pulse train signal to the pump controller, which then modulates the output of solid-state relays which gate power to each pump. Pumps are powered using a standard 12V/2A DC power supply, which is routed through a switch to control the pumps’ direction. While the pump controller is compatible with any pump that uses a standard DC motor (i.e. not a stepper motor, or a DC brushless motor), the aluminum pump plate that integrates the pump controller and the pumps is specifically designed for the Boxer 9QX pumps.

To control the pumps, we designed a pump control PCB (Figure 4B) which interfaces directly with the SpikeGadgets ECU and operates as a breakout board for controlling maze accessories (all digital outputs are buffered). DIO ports 1-8 are each connected to an RJ-45 port (pinout in Figure 4C), which we typically connect to the reward ports, but are fairly generic and can be used in a variety of ways. Ports 9-16, meanwhile, control the solid state relays used to gate power to the pumps, with filtering capacitors to protect the board from back EMF. By using standard JST-PH 2 connectors to connect to the peristaltic pumps, this board can be easily used to modulate power to a range of devices as necessary. Importantly, we have not observed electrical artifacts caused by the pumps on neural recordings made in close proximity to the PER (Figure 4-1). The pumps do make an audible noise when operating (see Movie 1). However, especially after maze acclimatization, subjects do not show any signs of attending to pump sounds from either their PER or neighboring PERs, such as interruptions in movement trajectories, extended pauses, or changes in orientation. To minimize any potential cross-maze sound impacts, or if the task itself required sound cues, isolation might be achieved by installing soundproof barriers between each maze.

Critically, in order to achieve consistent reward delivery of an appropriate amount of milk at an appropriate speed, we designed a variety of pulse trains rather than simply turning the pump on for a precise duration. This modulates the duty cycle, and results in a slower, more consistent milk delivery.

For an approximately 30µL ”small” reward we used 3 10ms pulses separated by 20ms (33% duty cycle). For an approximately 130µL ”large” reward we used 10 10ms pulses separated by 20ms. In addition to increasing consistency, this strategy is preferable to slowing the pumps by modulating their voltage, because it retains the ability to run pumps at full speed during milk loading and cleaning. All the pumps for each maze draw from a single 1L reservoir of milk, streamlining the milk loading process and reducing waste. In addition, it greatly increases the efficiency of the cleaning process. After daily behavioral training is complete, pump direction is reversed via a switch on the pump control PCB which allows us to collect the milk for overnight storage at 4*^◦^*C. We then fill the reservoir with cleaning solution, return the pumps to their default direction, flush the tubing until clean, run ethanol through the tubing to sanitize, and then let dry.

### 2.5. Drainage

In an attempt to reduce the number of refuse containers, gutters were made so that up to 3 reward ports could share a single waste container (Figure 1A, Figure 2B). Additionally, some containers could be shared between 2 gutters, or between 2 home ports, reducing the total number of refuse containers for all 4 mazes to 8. Gutters were made from 2” ABS tubing, cut lengthwise, and epoxied to a simple acrylic frame. It is critical to ensure that the tubing leading from the reward ports to the gutters maintains a sufficiently steep angle so that liquid does not pool and clog. Alternatively, longer tubes leading to shared refuse containers may also be used in place of gutters as long as the tubing angle is sufficiently steep.

### 2.6. Maze Assembly and Affordability

A guiding principle of our maze apparatus is affordability. By designing a maze environment that could be consistently assembled in-house from lasercut acrylic pieces, we avoided the high costs of custom manufacturing by commercial vendors such as Tap Plastics, Maze Engineers, Med Associates, or similar. In sum, the cost of materials for a single maze environment (acrylic, 8020, and hardware) came to approximately $1,000 and could be assembled in less than one day. In comparison, a comparable custom acrylic environment from a commercial vendor was quoted over $10,000. We use 3D printed reward wells instead of purchasing nosepokes from Med Associates or similar. These can be ordered from online 3D printing companies such as Protolabs or Shapeways for about $90 each or printed with entry-level 3D printers in labs or makerspaces for even lower cost (in our case, $10 in resin each on a lab Form3 printer). The PER includes a neural data acquisition system (SpikeGadgets) which is is a substantial investment for any lab doing neurophysiology, but could be optional for running behavior if another behavioral control system is used. The NUC computers are highly affordable ($400 each) compared to typical desktop computers. Peristaltic pumps, at approximately $140 each, are far more cost-effective than programmable syringe pumps (New Era Systems, $700 each) in addition to dramatically reducing cleaning time and effort. The overhead cameras are an exception to our budget design choices; the Allied Vision Manta cameras and lenses ($1,500 each) provide exceptional time synchronization abilities but could certainly be replaced with more affordable options if desired. Altogether, we estimate an up-front cost of approximately $15,000 for the complete 4-maze apparatus, command station, and all accessories.

### 3. Maze Task and Curriculum

The spatial memory task that we ultimately wanted subjects to perform is a challenging memory-dependent ”win-stay/lose-switch task”, related to prior work (Gillespie et al. 2021). Reliable performance is best achieved by using several simpler task variants (”steps”) to gradually shape behavior toward the final task (Holleman et al. 2019), which we refer to as our ”curriculum”. Each step of the curriculum is designed to build upon knowledge from previous steps, with advancement criteria designed to minimize both frustration and complacency in order to encourage engagement and effective learning. Well-defined advancement criteria enable automated advancement through the curriculum while also helping to standardize the learning process across subjects and cohorts. Breaking down the learning of the full task into simpler components is not only easier for the subject, but also opens up the possibility of studying the neural correlates of the learning process with more structure and specificity (Shujah et al. 2024) — thus both the training steps and the final task are scientifically valuable. Furthermore, by ensuring that all subjects meet standardized performance thresholds before advancing to the next step of the training process, this approach allows us to quantify measures of performance at each step, giving us valuable insight into the learning process. We focus on our 6-arm task and its training steps here, but many other types of tasks could be implemented, each with their own training curriculum, within this or other maze environments.

### 3.1. Task

For all 6-arm task steps, the subject must perform structured trials which consist of a nose poke at the home port, then a visit to a single arm of the maze, then a return to the home port (Figure 5A). Each trial begins when the subject triggers the illuminated home port and receives a small reward. Next, some number of the outer ports illuminate, and the subject must choose one to visit; receiving a large reward if the arm is a goal arm, and nothing otherwise. After visiting a single arm port the subject must return home to begin the next trial. The first few task steps are visually cued: rewarded outer arm ports are lit while unrewarded outer arm ports are not, so visiting any lit arm port will result in the subject receiving a reward (Figure 5B,D). Later, subjects transition to memory-dependent task steps, during which both rewarded and unrewarded ports are lit and subjects must discover and remember which one(s) deliver reward (Figure 5B,D). In later task steps, goal arm locations are changed in a ”block” structure (after 15 goal arm visits; Figure 5C). Within each goal block, the ”search phase” is the set of trials until the goal arm has been identified, and the ”repeat phase” includes all subsequent trials; the search phase includes the first rewarded trial. When the goal block is complete, new goal arm(s) are drawn, and the subject must begin a search phase to find the new goal arm(s). Any deviation from the described trial structure, such as going from one arm port to another, initiates a “lockout”, where all wells are de-illuminated/deactivated for a predetermined interval (typically 15s). After the lockout is over, the home well illuminates, and the subject must return home to start the next trial. Depending on the maze parameters, the epoch ends after a set number of trials, a set number of goal blocks, or a set amount of time (typically a maximum of 70 minutes from the epoch start). At this time, all wells are automatically de-illuminated/deactivated, and subjects are left to wait for the experimenter to remove them from the maze.

**Figure 5:**
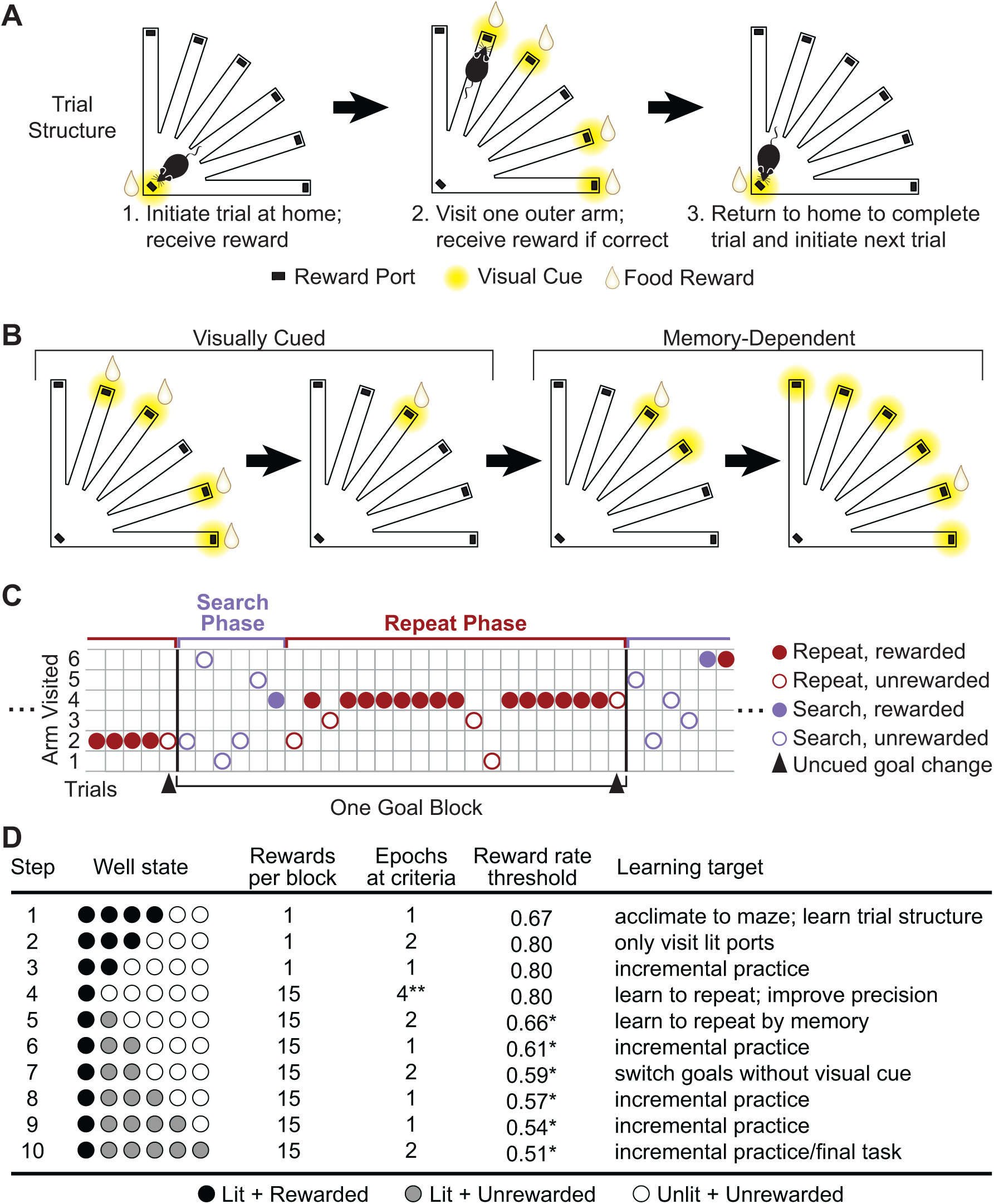
Training curriculum: A Visual schematic of a single example trial at step 1, including the initiating poke at the home port, choice of one outer arm, and the return to home to complete the trial. B Visual schematic of training progression, illustrating steps 1, 4, 5, and 10 of the training curriculum. C Example trial series for step 10 behavior, illustrating the classification of search and repeat trials. D A table giving an overview of the 10-step training curriculum. Outer reward ports exist in one of 3 states: Lit + Rewarded, Lit + Unrewarded, and Unlit + Unrewarded. In steps 5 and 6, when switching goal blocks, the previous goal well is never lit, giving the subjects a visual cue when it is time to switch goals. In steps 7 through 10, however, the previous goal well is always lit (unrewarded), forcing the subjects to learn to switch goals on their own. * indicates goal rate thresholds that were used for the female cohort and are recommended. The thresholds for these steps for the male cohort were marginally higher: 0.70, 0.66, 0.63, 0.60, 0.58, and 0.55, respectively. ** indicates the recommended number of epochs at step 4; only 1 was required for both the male and female cohorts.

### 3.2. Curriculum

Before beginning the 6-arm curriculum, subjects are first trained to trigger reward wells in order to receive milk reward on a simple linear track. Subjects must first earn >50 rewards in a 45-60 minute training session before advancing to the maze; this typically takes 3-5 training sessions.

The full 6-arm curriculum consists of 10 increasingly difficult steps as outlined in Figure 5D. Each step in the curriculum is characterized by a set of parameters: the number of goal ports, the number of lit ports, the number of rewards per goal block, the number of trials in the epoch, and the goal rate threshold (#rewarded trials / #trials over the entire epoch). The number of goal ports determines how many ports will be chosen as goals each block. The number of lit ports is the total number of reward ports with their white LEDs turned on during the outer arm phase of the trials. These ports are selected randomly once per block alongside the goal port, and exist as all options the subject must explore when searching for the goal (goal ports are always lit). Ports which are lit but unrewarded will never give a reward during the current goal block. The number of rewards per goal block determines how often the goal changes. Lastly, for an epoch to be considered “successful”, the subject must complete at least the minimum number of trials (typically 50 trials) while achieving a goal rate greater than or equal to the threshold goal rate. Goal rate thresholds for memory-dependent steps are generally calculated as 75% of the rate a perfectly-trained subject would achieve if they were maximally unlucky, minus an additional 5% buffer for the memory-based steps. At each step in the curriculum, subjects must complete a specified number of successful epochs before moving on to the next step. After the completion of step 10, subjects are considered fully trained, although additional epochs of behavior may be collected to fully characterize final performance levels.

### 3.3. Curriculum Implementation

Setting task parameters manually, especially with increasing cohort size and parallel training sessions, is extremely prone to experimenter error, while also being tedious, stressful, and time consuming for the experimenter. To counter this, each subject’s environment parameters and learning progress are stored in a text file in order to fully automate the subject’s progress through the curriculum. This text file is associated with the subject’s name, and is automatically read/updated by the same python script which controls the maze environment. In addition to being less error-prone, this strategy enforces uniform and objective environment parameter selection between training stages across subjects, removing any experimenter variability in deciding when to advance subjects to the next task step.

To safeguard against any errors which may occur during the epoch, the parameter file is updated as trials are completed, ensuring that it is as upto-date as possible at all times. Furthermore, a copy of the parameter file is also added to the behavior log at the beginning and end of each epoch, which makes the parameter set accessible for downstream data processing. This also preserves a record of which changes were made to the file by the maze code, and which changes were made by the experimenter between epochs (atypical, but occasionally necessary).

### 3.4. Arm Selection

Due to thigmotaxis, subjects tend to begin behavioral training with an innate bias towards arms 1 and 6, since those arms share a continuous wall with the home area. As subjects explore the maze and learn the task, they tend to develop idiosyncratic biases towards and against specific arms, which confounds behavioral analyses. To counter these biases, an arm selection algorithm (Appendix A) was implemented which tracks the subject’s arm visits, and draws from a probability distribution which favors less frequently visited arms, encouraging subjects to visit all arms as evenly as possible.

While simple in its aims, there were a few important considerations that we needed the algorithm to account for. First, if the algorithm ever pushes the probability of selecting a particular arm to effectively 0, subjects can and will over-correct, learning to completely avoid that arm, which is hard to recover from (in general, we found it easier to train subjects away from their favorite arms rather than towards arms they were averse to). To counter this, a lower bound was selected for the arm selection probability (typically 0.01 *−* 0.02).

Another issue we encountered was the algorithm being somewhat unresponsive, and failing to select arms which used to be a subject’s favorite long after the subject had learned to visit that arm less. While this could be countered by making the algorithm more aggressive, that comes at the cost of the algorithm becoming overly sensitive to the randomness inherent to goal selection and subjects’ behavior. So, to increase responsiveness of the algorithm, we added a parameter which weights recent arm visits more heavily than older arm visits; allowing subjects’ past behavior to quickly become completely irrelevant, while also only responding to new behaviors after a certain level of consistency is demonstrated. Two additional parameters were included which control whether the rewarded visits or unrewarded visits are more influential, and how aggressively the algorithm attempts to push subjects towards even exploration.

With these four parameters, our arm selection algorithm is simple, flexible, and responsive, while avoiding some of the pitfalls which can seriously hamper learning.

### 3.5. Data Outputs

For post-hoc analysis, we need reliable records of all behavioral events, such as port triggers, port illumination state, and reward delivery times. Further, we need a system that is resilient to common experimenter errors such as typos in filenames and recording mistakes. We developed a system of multiple logs, each with varying levels of detail, which are initialized at the start of each epoch and which automatically close themselves at the end of the epoch.

The first is a highly detailed log containing the maze parameters, initialization statements, and all maze events. Furthermore, it is preemptively segmented into goal blocks to make data processing as easy as possible. The second log is much more compact, containing one line for the maze parameters and a table of important trial events. Each trial contains the data for the home visit, the outer arm visit, goal block details, and the details for any lockout port visits. The data recorded for each port visit includes the time and duration of the IR beam break, which port was triggered, and whether or not a reward was delivered. While not as detailed as the full log, this format is extremely easy to process, and requires no detective work to assemble the trial landmarks, making it very convenient for data processing.

Additionally, each subject’s arm visits are plotted live at the control station, meaning that the experimenter can monitor all subjects’ progress simultaneously (Figure 1-1B). This both allows the experimenter to gain a deeper understanding of how subjects are performing the task without having to constantly watch the subjects and makes it easy to quickly detect any maze malfunctions. The videos collected by overhead cameras during behavior are also saved for post-hoc position tracking or pose estimation as needed (Movie 2, 3).

### 3.6. Resources and Code Accessibility

As many aspects of this maze apparatus have been made open-source as possible; links to solid models, laser cutter drawings, and PCB design files can all be found in Appendix B, along with a complete BOM. Additionally, the python codebase used for task logic, performance tracking, and curriculum advancement is freely accessible (Saathoff and Gillespie 2025a,b) without restrictions.

### 3.7. Experimental Model and Subject Details

To test the automated, parallelized maze apparatus, we trained cohorts of 8 male and 8 female Fischer x Brown Norway F1 rats from the NIA colony (Charles River). Each cohort included 4 young (4-9 months) and 4 aged (28-32 months) subjects. All animals were kept on a 12 hr light-dark cycle (lights on 7am-7pm) with *ad libitum* access to food (standard rat chow) in a temperature-and humidity-controlled facility. Beginning one week prior to training, rats were singly housed and food restricted to 85% of their free-feeding weight. Data were collected from over 32 weekdays (male cohort) and 37 weekdays (female cohort). Food restriction was maintained on weekends, but behavioral training was not administered. Each subject completed 2-3 epochs per training day. Subjects in each cohort were divided into two subgroups of 4 each, each containing two young rats and two aged rats; the subgroup which ran first in the day was alternated each day. Each subject was assigned to a specific maze, with one young rat and one aged rat assigned to each maze. Behavioral training for the male and female cohort was conducted serially, not interleaved, with thorough cleaning of the room between cohorts to avoid any overlap in odor cues. All procedures were approved by the University of Washington Institutional Animal Care and Use Committee.

## 4. Results

The male cohort was trained using the curriculum shown in Figure 5D, with minor differences indicated by asterisks and detailed in the figure caption. For this cohort, subjects received a small reward at the home port and a large reward at outer arm ports. Subjects showed a rapid increase in reward rate during the visually cued epochs, all reaching performance levels well above chance, but also demonstrating substantial individual variability in learning rate and ultimate performance level (Figure 6A). The memory-dependent version of the task was learned much more slowly, but ultimately all subjects reached performance rates well above chance (Figure 6B). Two of our aged subjects failed to reach the set criterion for memory-dependent step 8 after many epochs (10 epochs for A1, and 8 epochs for A3) and were manually advanced to step 9 for a single epoch, and then to step 10. Despite this, we found no obvious differences in performance at the final task between aged subjects who completed the curriculum, and aged subjects who were advanced manually. Additionally, we found no significant difference between age groups in repeat phase reward rate (Figure 6C) or the length of the search phase; a measure indicative of search efficiency (Figure 6D). The amount of training required to reach criterion for most steps was suggestive of slower learning in the aged subjects, particularly for the first few steps of the memory-dependent task variant (Figure 6E,G), although this also did not reach statistical significance likely due to our small sample size. The most drastic difference between age groups was in trial duration (Figure 6F), as aged subjects moved far more slowly than young subjects. Despite their slow speed, aged subjects generally completed the designated number of trials or goal blocks and only rarely timed out.

**Figure 6:**
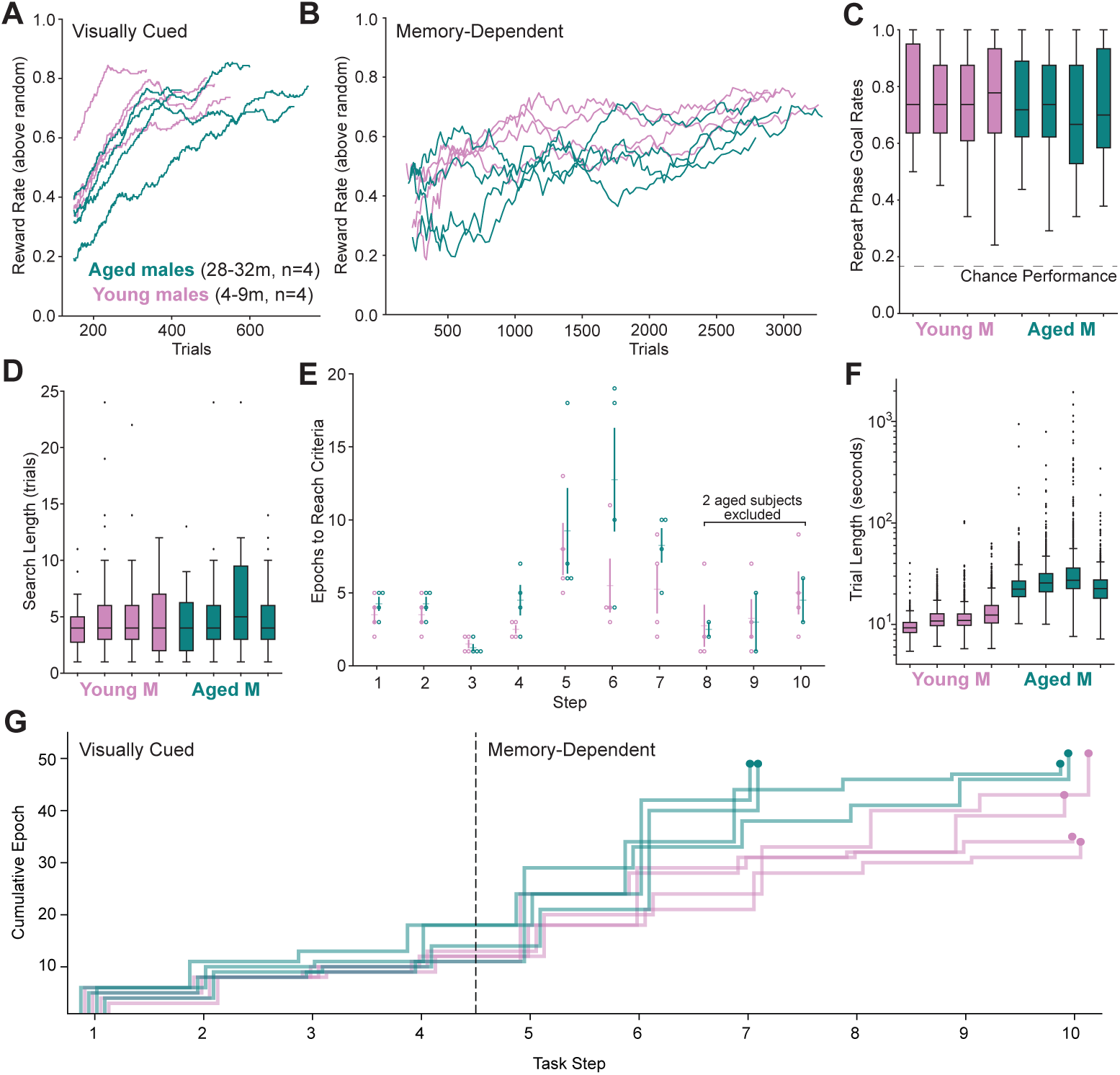
Male Cohort Results: A Learning curves by subject for the visually cued task steps. This plot shows the reward rate for each subject minus the reward rate a random agent would achieve, linearly-normalized such that 0 is the reward rate of a random agent, and 1 is the reward rate a subject would achieve with a 100% goal rate. Data was smoothed with a 150 trial moving average. n*_trials_* = [Y 1 : 510, Y 2 : 550, Y 3 : 336, Y 4 : 490, A1 : 600, A2 : 424, A3 : 713, A4 : 750]. B Learning curves by subject for the memory dependent task steps, linearly-normalized such that 0 is the reward rate of a random agent (that only selects lit wells), and 1 is the reward rate a subject would achieve with a 100% goal rate. An overall reward rate was calculated for each goal block and then smoothed with a moving average of 10 goal blocks (this eliminates oscillation due to the search phase/repeat phase structure of the task). n*_trials_* = [Y 1 : 3303, Y 2 : 3096, Y 3 : 3498, Y 4 : 3172, A1 : 3206, A2 : 3537, A3 : 3044, A4 : 2990] (includes plateaued behavior). C Median reward rate after finding the goal (repeat phase) during ”fully trained” memory-dependent epochs, after criterion was met for step 10. Effect of age evaluated by linear mixed-effect model (Python statsmodels mixedlm function): p=0.205, n*_blocks_* = [Y 1 : 36, Y 2 : 70, Y 3 : 54, Y 4 : 68, A1 : 32, A2 : 35, A3 : 31, A4 : 39] (p value calculated using Powell’s Method, instead of the default BFGS optimization method for the Mixed Effects Linear Regression). D Median length of search phase per goal block during ”fully trained” memory-dependent epochs, after criterion was met for step 10. Effect of age evaluated by linear mixed-effect model: p=0.123, n*_blocks_* = [Y 1 : 36, Y 2 : 70, Y 3 : 54, Y 4 : 68, A1 : 32, A2 : 35, A3 : 31, A4 : 39]. E Average number of training epochs required to meet progression criteria for each step; subjects excluded for steps they were unable to complete (see panel G). Ranksum results by step (Appendix C) do not demonstrate a significant effect of age on learning rates for this task in this small cohort of males. F Median length of time for each trial (time from initial home poke to the home poke initiating the next trial). Effect of age evaluated by linear mixed-effect model: p < 0.001, n*_trials_* = [Y 1 : 848, Y 2 : 1711, Y 3 : 1358, Y 4 : 1672, A1 : 781, A2 : 961, A3 : 909, A4 : 1010]. G Step plot showing the cumulative advancement of each male subject through the curriculum. Note that horizontal offsets are added so each individual subject line is visible. The circle at the end of each line indicates the furthest epoch completed by that subject prior to manual advancement.

The female cohort was also trained using the curriculum shown in Figure 5D, with only one minor difference noted in the figure caption. Importantly, however, this cohort was provided only a small reward at both the home and outer arm ports, since the female rats were much smaller and more likely to satiate with the amount of milk reward provided to the male cohort. Perhaps as a result of this difference, the female cohort learned both the visually cued and memory-dependent task steps more slowly (Figure 7A,B). Further, three aged subjects and one young subject were unable to meet the performance criteria required to advance through the full curriculum (Figure 7E,G). These subjects were manually advanced to the final step 10 task variant, where we saw that aged subjects had a significantly lower reward rate during repeat phase (Figure 7C) and trended toward longer search phases (indicative of inefficient searching; Figure 7D). However, these results could reflect the altered relative reward amounts and/or the manual advancement rather than reflecting true differences in performance ability. Prior to becoming unable to advance, however, aged subjects showed a trend toward slower learning, especially in the memory-dependent steps, than the young subjects (Figure 7B,E,G). They also moved far more slowly (Figure 7F). Both of these results are consistent with the results of the male cohort.

**Figure 7:**
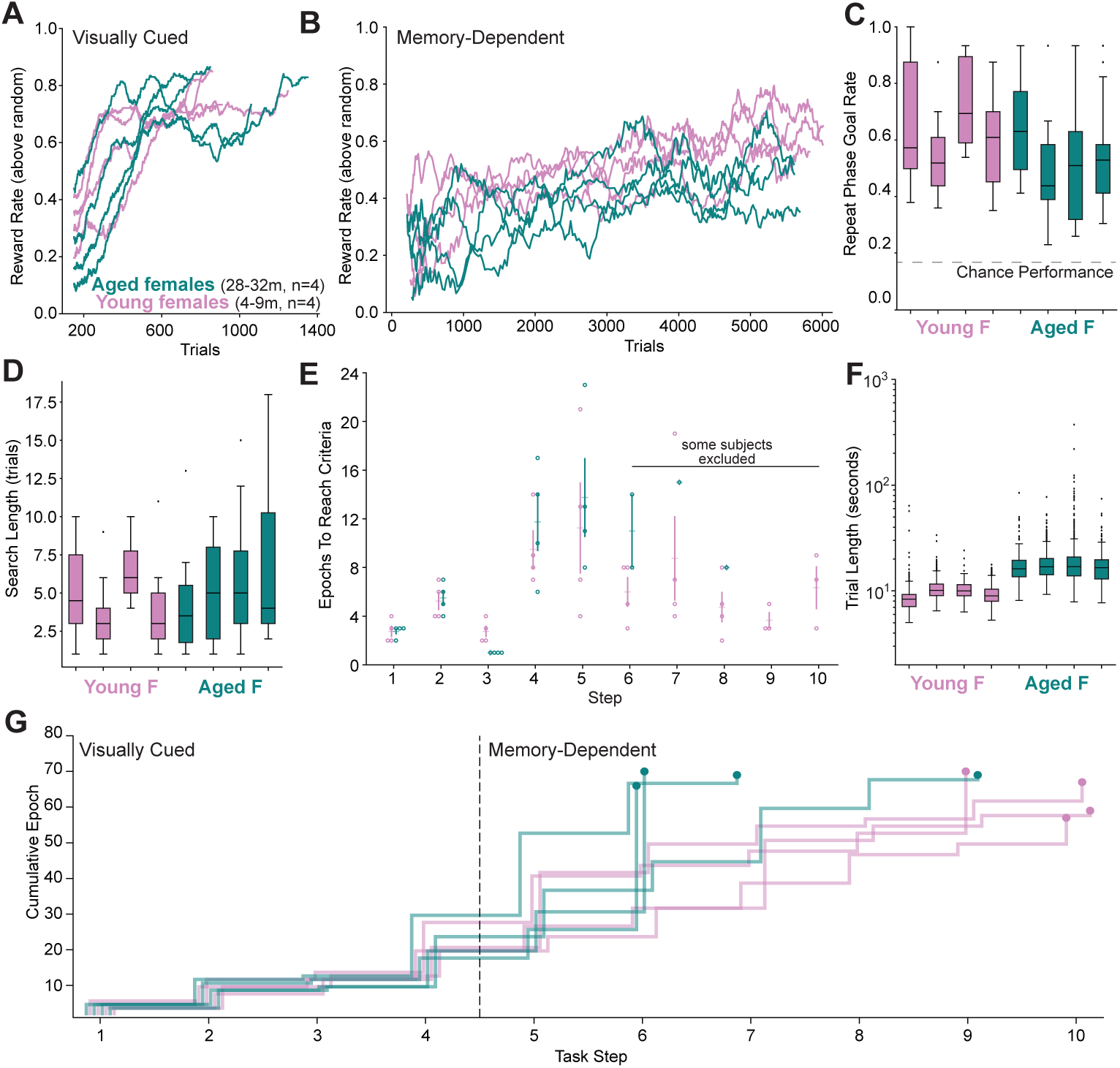
Female Cohort Results: A Learning curves by subject for the visually cued task steps. This plot shows the reward rate for each subject minus the reward rate a random agent would achieve, linearly-normalized such that 0 is the reward rate of a random agent, and 1 is the reward rate a subject would achieve with a 100% goal rate. Data was smoothed with a 150 trial moving average. n*_trials_* = [Y 1 : 1250, Y 2 : 910, Y 3 : 860, Y 4 : 840, A1 : 1060, A2 : 750, A3 : 850, A4 : 1350]. B Learning curves by subject for the memory dependent task steps, linearly-normalized such that 0 is the reward rate of a random agent (that only selects lit wells), and 1 is the reward rate a subject would achieve with a 100% goal rate. An overall reward rate was calculated for each goal block and then smoothed with a moving average of 10 goal blocks (this eliminates oscillation due to the search phase/repeat phase structure of the task). n*_trials_* = [Y 1 : 5675, Y 2 : 5975, Y 3 : 6135, Y 4 : 6175, A1 : 5765, A2 : 5800, A3 : 5880, A4 : 5280] (includes plateaued behavior). C Median reward rate after finding the goal (repeat phase) during ”fully trained” memory-dependent epochs, after criterion was met for step 10. Effect of age evaluated by linear mixed-effect model (Python statsmodels mixedlm function): p=0.046, n*_blocks_* = [Y 1 : 12, Y 2 : 13, Y 3 : 6, Y 4 : 17, A1 : 8, A2 : 17, A3 : 18, A4 : 20] (p value calculated using Powell’s Method, instead of the default BFGS optimization method for the Mixed Effects Linear Regression). D Median length of search phase per goal block during ”fully trained” memory-dependent epochs, after criterion was met for step 10. Effect of age evaluated by linear mixed-effect model: p=0.097, n*_blocks_* = [Y 1 : 12, Y 2 : 13, Y 3 : 6, Y 4 : 17, A1 : 8, A2 : 17, A3 : 18, A4 : 20]. E Average number of training epochs required to meet progression criteria for each step; subjects excluded for steps they were unable to complete (see panel G). Ranksum results for steps completed by at least 2 aged female subjects (Appendix D). Most steps did not demonstrate a significant effect of age on learning rates in this small female cohort, although aged subjects advanced through step 3 slightly but significantly more quickly than young subjects. F Median length of time for each trial (time from initial home poke to the home poke initiating the next trial). Effect of age evaluated by linear mixed-effect model: p < 0.001, n*_trials_* = [Y 1 : 420, Y 2 : 490, Y 3 : 210, Y 4 : 630, A1 : 280, A2 : 910, A3 : 910, A4 : 910] G Step plot showing the cumulative advancement of each female subject through the curriculum. Note that horizontal offsets are added so each individual subject line is visible. The circle at the end of each line indicates the furthest epoch completed by that subject prior to manual advancement.

## 5. Discussion

The results from the pilot cohorts demonstrate that the mazes perform largely as expected, achieving our major goals of training a cohort of rats with minimal experimenter interference on a complex curriculum. In contrast to prior training regimes for a similar task (Gillespie et al. 2021), our apparatus allowed us to train twice as many rats in about half the time. Further, our training process now allows us to capture different learning phases in a regimented and comparable manner. The gradual advancement through increasingly complex steps allowed more rats to continue participating, rather than being dropped from the cohort due to poor motivation or low performance. While aged subjects were certainly capable of learning and performing the final memory-dependent task variant, differences in movement speed and learning rate suggest that this is an appropriate task in which to study the impact of age on memory processes. Additionally, with experimenter-induced variability minimized, the individual variability remaining in both age groups further suggests that this task may be well suited to explore the neural basis of individual variability in cognitive performance.

Two aspects of the behavioral analysis were particularly illuminating. First, the dramatic difference in movement speed, and thus trial length, between the age groups for both sexes was striking. While unsurprising (Birren and Fisher 1995; McQuail and Nicolle 2015), this underscored the importance of having each epoch be limited by trial or goal block criteria, rather than by time (as is more common). This ensured that the aged and young subjects received similar amounts of experience with each epoch and will be essential for any studies of aged subjects. It was also encouraging that, when given sufficient time, aged subjects were motivated and able to engage with the task similarly to the young subjects. Second, the implementation of such a gradual curriculum was effective, as indicated by the lack of any stark performance dropoff caused by progression to a specific task step and the continued engagement of every subject throughout the curriculum, even for subjects who did not achieve all advancement criterion. Further, the inclusion of learning both a visually cued and memory-dependent task separates the learning of distinct task rules, allowing for future investigation into the neural correlates of each rule separately.

### 5.1. Limitations

While very infrequent, hardware errors and experimenter errors did still occasionally occur. Hardware errors tended to be reward port failure due to beam break sensors failing or tubing leaks causing failure in reward delivery. These issues were difficult to catch by inspection of the apparatus, but became readily apparent during behavior as subjects would rapidly learn to avoid ports that never delivered reward and these biases were clearly visible on the live performance plots. Watching for these issues required attentiveness from the experimenter, but once caught were promptly fixed, and epochs were re-done as necessary (dropping any data which was adversely affected). Experimenter error tended to be minor, and extra epochs were performed to replace impacted data.

A second limitation that our pilot cohorts revealed was the impact of setting advancement criteria at a level achievable by all subjects, while also enforcing sufficient learning of the current task variant. The two aged males who were unable to advance past step 8 were still performing well above chance, and nearly at criterion — suggesting not that they failed to learn, but that the criterion was set unreasonably high. Furthermore, after manual advancement, their final performance was not different than the other aged subjects. This strongly suggests that the issue was in fact one of parameter choice rather than a true performance difference. In the female cohort, we lowered the advancement criteria for memory-dependent steps but also introduced another change: we reduced the amount of reward provided at outer arm ports. We found that more female rats were unable to advance: three aged subjects and one young subject. We hypothesize that the reduction in reward amount may have substantially reduced motivation as the task variants became more difficult, obscuring any benefit of the lower advancement criterion. Another possibility is that lowering the outer reward amount caused subjects to devalue the outer arms and rely on the home well to provide reward, without learning to choose the outer arm optimally. In future behavioral studies, we will retain relatively larger reward amounts at the outer arm ports and we may further refine the advancement criteria, potentially by incorporating metrics other than or in addition to simple overall reward rate. Future implementation of this task should give careful consideration to advancement criteria, especially considering the specific goals of the experiment. For instance, fixed advancement criteria may be essential for comparing learning between experimental cohorts, while more flexible criteria may be appropriate if the goal is to train all subjects to perform a final high-priority task variant. Since many factors (rat strain, age, experimental intervention, room setup, experimenter style, and more) may influence behavioral performance, more pilot training cohorts will be essential to inform performance expectations when developing new tasks or curricula.

### 5.2. Future Outlook

While the current apparatus substantially extends our ability to train and collect data on larger cohorts of animals, it is still far from truly high-throughput behavior as other groups have achieved (Hao et al. 2021; Poddar et al. 2013). One major difference is our continued use of discrete, spaced training sessions in specific maze environments separate from the home cage, which will always require more time and experimenter involvement than incage, unsupervised, continuous engagement tasks. Another key limitation to higher throughput is the requirement for overhead camera-based tracking throughout the entire maze. If position information were unnecessary, or could be collected via alternate means (via inertial sensors (Fayat et al. 2025), for instance), then it would be feasible to stack 3-4 mazes vertically and run 12-16 subjects simultaneously with the same spatial footprint. Some updates to drainage and cleaning strategies would be needed in this case, but all task control and logging software would seamlessly extend to the stacked use case. Such stacking of the mazes could allow comparable throughput to other highly successful behavioral training approaches (Hao et al. 2021; Poddar et al. 2013), and would be the first to our knowledge to achieve this level of throughput for a spatial navigation task.

Even at its current capacity, the maze apparatus opens up exciting possibilities for efficiently testing larger cohorts of male and female subjects, rodent models of disease, and performing longitudinal behavioral assessment of cohorts across the lifespan — all of which will be fascinating comparators to the expansion of our existing pilot study of age-related behavioral differences. Since the apparatus is also fully compatible with wireless neural recordings and overhead position tracking, future studies will include the assessment of neural activity patterns that underlie the learning and memory processes that we set out to study. The ability to test larger cohorts of animals with minimal pre-selection will allow us to capture and characterize performance across a much wider range of abilities, facilitating our understanding of the neural basis of individual performance differences. Another key challenge in studying individual behavioral differences is the need to identify biologically-driven, rather than experimentally-induced, variability. The automation and parallelization of the current maze apparatus suggests that the variability we see in pilot cohort performance may reflect, and allow the study of, true individual differences.

We hope that the low cost and open source nature of many aspects of our task apparatus will make it a useful resource for others striving to scale up and increase the automation of their behavioral testing approaches. In the described maze environment, the flexible nature of the task logic can allow for many different tasks to be implemented, such as sequence learning tasks, alternation tasks (Frank et al. 2000; Kastner et al. 2020), or even pure foraging tasks. The use of alternative maze environments equipped with our, or other, maze features (Porter et al. 2025) opens endless possibilities for structured, efficient behavioral testing. Regardless of behavioral paradigm, a shift towards parallelized training offers improved control of other influences on behavior, such as circadian rhythms, in addition to increased efficiency. Finally, the adoption of a structured curriculum with trial-based advancement criteria, rather than time-based advancement criteria, will be beneficial for accurately and rigorously capturing behavioral differences across groups of subjects no matter the species or task.

## 6. Author Credit Statement

A.K.G. conceived the study, acquired funding, and supervised the project. V.T.S. and C.I.O. designed the experimental apparatus. V.T.S., C.I.O., and A.K.G. built the experimental apparatus. V.T.S. wrote behavioral and analysis software. V.T.S, J.A.Q., and Z.D.P. collected data. A.K.G. and V.T.S. designed the curriculum. V.T.S. and J.A.Q. analyzed the data with input from A.K.G. V.T.S., J.A.Q., and A.K.G. prepared data visualizations and wrote the manuscript with input from all authors.

## 7.#Acknowledgements

The authors are grateful to all Gillespie Lab members for their help constructing the maze apparatus and their insightful feedback on the manuscript. This work was supported by the NIH Brain Initiative/NIA (grant R00AG068425 to A.K.G.), the Simons Collaboration on Plasticity and the Aging Brain (grant 1169066 to A.K.G.), and the Esther A. & Joseph Klingenstein Fund (Fellowship Award in Neuroscience to A.K.G.).

## Appendix A. Arm Selection Algorithm

For most of training, the following algorithm was used to track subject’s arm visits, and select which arms to use as the goal arms. The premise is to maintain a value for each arm which quantifies how much the subject has visited that arm recently compared to other arms; these values are updated every time the subject visits as well. Then, when it is time to select a goal, the array of values is passed though an algorithm to generate a probability distribution where the most visited arms are the least likely to be selected, and the least visited arms are the most likely to be selected. Goals are then drawn from this distribution as needed.

There are four parameters which control the algorithm:

- α : weights between rewarded visits and unrewarded visits. α *∈* (0, 1); should be close to 0.5
- β : a ”temperature” parameter which controls how aggressively the algorithm will push subjects to explore all arms evenly. β < 0; shouldn’t be too large (i.e. β > *−*2 is recommended
- δ : a floor on how unlikely any particular arm can become -this helps prevent subjects from learning to never visit an arm that used to be their favorite, creating a new arm they’re averse to. δ *∈* [0, 1/6); should be small (i.e. 0 < δ < 0.03)
- γ : a decay parameter which causes more recent data to be more important than older data. This decay is applied approximately once per epoch. γ *∈* (0, 1); should generally be larger than 0.5

Two of these parameters have derived quantities, which are useful in practice:

- 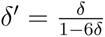
- 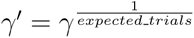

### Appendix A.1. Weighted Visits Update Function

well is the index of the well the subject just poked, and reward is either a 0 or 1 based on whether the subject received a reward. All vector operations are element-wise, and the 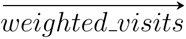 array is updated every time a subject visits an outer arm.

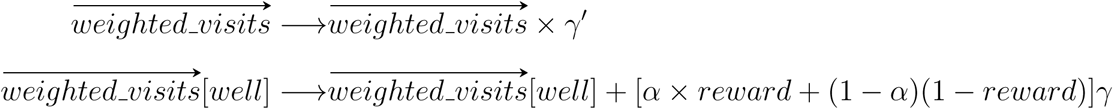

### Appendix A.2. Probability Distribution Function

Arms are chosen at random based on a probability distribution from the 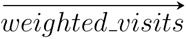 array. The probability distribution is re-computed using the formula below every time new goals need to be selected (i.e. at the start of every goal block).

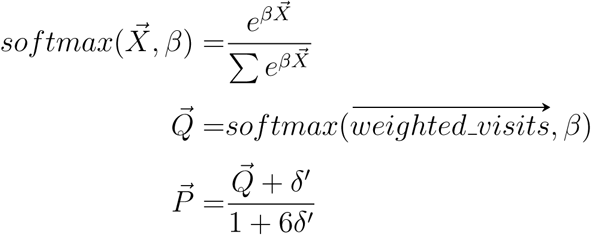

### Appendix A.3. Goal Selection Notes

- When only 1 goal is being selected (steps 4+ of the curriculum), care is taken to ensure that no arm is ever selected as the goal arm twice in a row.
- When subjects are searching for a single goal arm among several lit options, the goal selection algorithm presented above was found to decrease task engagement more than it helped minimize subjects’ search biases. So, after step 6 of the curriculum, goals are selected at random from a uniform distribution in such a way that random selection biases are eliminated. To do this, the next 6 goals are selected at a time such that each arm will be chosen once over the next 6 goal blocks. This information is stored in the parameter file to ensure that this arm selection persists between epochs. This is necessary because the number of goals drawn for each subject is relatively small, so random fluctuations in goal selection can introduce biases into the subjects’ behavior. When re-drawing goals, care is taken to ensure that no arm is selected as the goal arm twice in a row (the initial shuffling process prevents this within a set of 6 goals, but a check/reshuffle is necessary between sets of 6 goals).

## Appendix B. Resources

- Maze Design: https://doi.org/10.5281/zenodo.19701427
- Reward Port Design: https://doi.org/10.5281/zenodo.19711190
- Pump Plate Design: https://doi.org/10.5281/zenodo.19711326
- Bill of Materials: https://doi.org/10.5281/zenodo.19699417
- Maze Code: https://doi.org/10.5281/zenodo.19699679

## Appendix C. Statistics for Figure 6E, Male Ranksum Results

| step | 1 | 2 | 3 | 4 | 5 | 6 | 7 | 8 | 9 | 10 |
| --- | --- | --- | --- | --- | --- | --- | --- | --- | --- | --- |
| $n_{young}$ | 4 | 4 | 4 | 4 | 4 | 4 | 4 | 4 | 4 | 4 |
| $n_{aged}$ | 4 | 4 | 4 | 4 | 4 | 4 | 4 | 2 | 2 | 2 |
| ranksum statistic | -0.866 | -0.866 | 0.577 | -1.44 | -0.289 | -1.44 | -1.44 | -0.694 | 0.231 | 0.00 |
| p value | 0.39 | 0.38 | 0.56 | 0.15 | 0.77 | 0.15 | 0.15 | 0.49 | 0.82 | 1.0 |

## Appendix D. Statistics for Figure 7E, Female Ranksum Results

| step | 1 | 2 | 3 | 4 | 5 | 6 | 7 | 8 | 9 | 10 |
| --- | --- | --- | --- | --- | --- | --- | --- | --- | --- | --- |
| $n_{young}$ | 4 | 4 | 4 | 4 | 4 | 4 | 4 | 4 | 4 | 4 |
| $n_{aged}$ | 4 | 4 | 4 | 4 | 4 | 2 | 1 | 1 | 0 | 0 |
| ranksum statistic | -0.144 | -0.289 | 2.309 | -0.722 | -0.722 | -1.389 | - | - | - | - |
| p value | 0.885 | 0.773 | 0.021 | 0.470 | 0.470 | 0.165 | - | - | - | - |

## References

Astur, Robert S., et al. (Apr. 2002). “Humans with hippocampus damage display severe spatial memory impairments in a virtual Morris water task”. eng. In: Behavioural Brain Research 132.1, pp. 77–84. issn: 0166-4328. doi: 10.1016/s0166-4328(01)00399-0.

*Bcontrol* (2025). url: https://brodylabwiki.princeton.edu/bcontrol/index.php/Main_Page (visited on 11/01/2025).

Birren, J. E. and L. M. Fisher (1995). “Aging and speed of behavior: possible consequences for psychological functioning”. eng. In: Annual Review of Psychology 46, pp. 329–353. issn: 0066-4308. doi: 10.1146/annurev. ps.46.020195.001553.

Chesler, Elissa J., et al. (Nov. 2002). “Influences of laboratory environment on behavior”. en. In: Nature Neuroscience 5.11. Publisher: Nature Publishing Group, pp. 1101–1102. issn: 1546-1726. doi: 10.1038/nn1102-1101. url: https://www.nature.com/articles/nn1102-1101 (visited on 11/01/2025).

D’Hooge, Rudi and Peter P De Deyn (Aug. 2001). “Applications of the Morris water maze in the study of learning and memory”. In: Brain Research Reviews 36.1, pp. 60–90. issn: 0165-0173. doi: 10.1016/S0165-0173(01)00067-4. url: https://www.sciencedirect.com/science/article/pii/S0165017301000674 (visited on 07/21/2025).

Fayat, Romain, et al. (Oct. 2025). “DISSeCT: An unsupervised framework for high-resolution mapping of rodent behavior using inertial sensors”. en. In: PLOS Biology 23.10. Publisher: Public Library of Science, e3003431. issn: 1545-7885. doi: 10.1371/journal.pbio.3003431. url: https://journals.plos.org/plosbiology/article?id=10.1371/journal.pbio.3003431 (visited on 11/01/2025).

Findlay, Graham, et al. (Mar. 2020). “The evolving view of replay and its functions in wake and sleep”. en. In: SLEEP Advances 1.1, zpab002. issn: 2632-5012. doi: 10.1093/sleepadvances/zpab002. url: https://academic.oup.com/sleepadvances/article/doi/10.1093/sleepadvances/zpab002/6124581 (visited on 03/11/2025).

Frank, Loren M. et al. (July 2000). “Trajectory Encoding in the Hippocampus and Entorhinal Cortex”. English. In: Neuron 27.1. Publisher: Elsevier, pp. 169–178. issn: 0896-6273. doi: 10.1016/S0896-6273(00)00018-0. url: https://www.cell.com/neuron/abstract/S0896-6273(00)00018-0 (visited on 07/21/2025).

Gillespie, Anna, et al. (Oct. 2021). “Hippocampal replay reflects specific past experiences rather than a plan for subsequent choice”. eng. In: Neuron 109.19, 3149–3163.e6. issn: 1097-4199. doi: 10.1016/j.neuron.2021.07.029.

Hao, Yaoyao et al. (May 2021). “Fully autonomous mouse behavioral and optogenetic experiments in home-cage”. In: eLife 10. Ed. by Brice Bathellier et al. Publisher: eLife Sciences Publications, Ltd, e66112. issn: 2050-084X. doi: 10.7554/eLife.66112. url: https://doi.org/10.7554/eLife.66112 (visited on 03/04/2025).

Holleman, Esther, et al. (Aug. 2019). “An incremental training method with automated, extendable maze for training spatial behavioral tasks in rodents”. en. In: Scientific Reports 9.1, p. 12589. issn: 2045-2322. doi: 10.1038/s41598-019-48965-w. URL: https://doi.org/10.1038/s41598-019-48965-w (visited on 07/21/2025).

Kastner, David B., et al. (Sept. 2020). “Memory Alone Does Not Account for the Way Rats Learn a Simple Spatial Alternation Task”. eng. In: The Journal of Neuroscience: The Official Journal of the Society for Neuroscience 40.38, pp. 7311–7317. issn: 1529-2401. doi: 10.1523/JNEUROSCI.0972-20.2020.

Lopes, Goņcalo and Patricia Monteiro (Mar. 2021). “New Open-Source Tools: Using Bonsai for Behavioral Tracking and Closed-Loop Experiments”. English. In: Frontiers in Behavioral Neuroscience 15. Publisher: Frontiers. issn: 1662-5153. doi: 10.3389/fnbeh.2021.647640. url: https://www.frontiersin.org/journals/behavioral-neuroscience/articles/ 10.3389/fnbeh.2021.647640/full (visited on 11/01/2025).

McQuail, Joseph A. and Michelle M. Nicolle (2015). “Spatial reference memory in normal aging Fischer 344 × Brown Norway F1 hybrid rats”. In: Neurobiology of Aging 36.1. Place: Netherlands Publisher: Elsevier Science, pp. 323–333. issn: 0197-4580. doi: 10.1016/j.neurobiolaging. 2014.06.030.

Morris, Richard G. M. (May 1981). “Spatial localization does not require the presence of local cues”. In: Learning and Motivation 12.2, pp. 239–260. issn: 0023-9690. doi: 10.1016/0023-9690(81)90020-5. url: https: //www.sciencedirect.com/science/article/pii/0023969081900205 (visited on 03/04/2025).

Moser, May-Britt, et al. (Feb. 2015). “Place Cells, Grid Cells, and Memory”. en. In: Cold Spring Harbor Perspectives in Biology 7.2, a021808. issn: 1943-0264. doi: 10.1101/cshperspect.a021808. url: http://cshperspectives.cshlp.org/lookup/doi/10.1101/cshperspect.a021808 (visited on 03/11/2025).

O’Keefe, J. and J. Dostrovsky (Nov. 1971). “The hippocampus as a spatial map. Preliminary evidence from unit activity in the freely-moving rat”. en. In: Brain Research 34.1, pp. 171–175. issn: 00068993. doi: 10.1016/0006-8993(71)90358-1. url: https://linkinghub.elsevier.com/retrieve/pii/0006899371903581 (visited on 03/11/2025).

Olton, David S. and Robert J. Samuelson (1976). “Remembrance of places passed: Spatial memory in rats”. In: Journal of Experimental Psychology: Animal Behavior Processes 2.2. Place: US Publisher: American Psychological Association, pp. 97–116. issn: 1939-2184. doi: 10.1037/0097-7403.2.2.97.

Pioli, Elsa Y., et al. (Mar. 2014). “An automated maze task for assessing hippocampus-sensitive memory in mice”. In: Behavioural Brain Research 261, pp. 249–257. issn: 0166-4328. doi: 10.1016/j.bbr.2013.12.009. url: https://www.sciencedirect.com/science/article/pii/S0166432813007523 (visited on 07/22/2025).

Poddar, Rajesh, et al. (Dec. 2013). “A Fully Automated High-Throughput Training System for Rodents”. en. In: PLOS ONE 8.12. Publisher: Public Library of Science, e83171. issn: 1932-6203. doi: 10.1371/journal.pone.0083171. url: https://journals.plos.org/plosone/article?id=10.1371/journal.pone.0083171 (visited on 07/21/2025).

Porter, Blake S. et al. (July 2025). “Adapt-A-Maze: An Open-Source Adaptable and Automated Rodent Behavior Maze System”. en. In: eNeuro 12.7. Publisher: Society for Neuroscience Section: Research Article: Methods/New Tools. issn: 2373-2822. doi: 10.1523/ENEURO.0138-25.2025. url: https://www.eneuro.org/content/12/7/ENEURO.0138-25.2025 (visited on 11/01/2025).

Qiao, Mu et al. (Feb. 2019). Mouse Academy: high-throughput automated training and trial-by-trial behavioral analysis during learning. en. Pages: 467878 Section: New Results. doi: 10.1101/467878. url: https://www. biorxiv.org/content/10.1101/467878v2 (visited on 11/01/2025).

Roumis, Demetris (2020). “Rhythmic Action Synchronizes Memory Replay During Reinforcement Learning”. en. PhD thesis. UCSF. url: https://escholarship.org/uc/item/10s205q3 (visited on 07/21/2025).

Saathoff, Violet and Anna Gillespie (July 2025a). gl-behavior. Version 1.0.0. — (July 2025b). gl-behavior-modules. Version 1.0.0.

Shujah, Maria, et al. (Sept. 2024). *Curriculum Learning: sequential acquisition of task complexity enhances neuronal discriminability*. en. Pages: 2024.09.10.612027 Section: New Results. doi: 10.1101/2024.09.10.612027. url: https://www.biorxiv.org/content/10.1101/2024.09.10.612027v1 (visited on 11/01/2025).

Siegle, Joshua H et al. (June 2017). “Open Ephys: an open-source, plugin-based platform for multichannel electrophysiology”. en. In: Journal of Neural Engineering 14.4. Publisher: IOP Publishing, p. 045003. issn: 1741-2552. doi: 10.1088/1741-2552/aa5eea. url: https://doi.org/10.1088/1741-2552/aa5eea (visited on 11/01/2025).

Steiner, Adam P. and A. David Redish (July 2014). “Behavioral and neurophysiological correlates of regret in rat decision-making on a neuroeconomic task”. eng. In: Nature Neuroscience 17.7, pp. 995–1002. issn: 1546-1726. doi: 10.1038/nn.3740.

Tolman, Edward C. (1948). “Cognitive maps in rats and men”. In: Psychological Review 55.4. Place: US Publisher: American Psychological Association, pp. 189–208. issn: 1939-1471. doi: 10.1037/h0061626.

Vorhees, Charles V. and Michael T. Williams (2014). “Value of water mazes for assessing spatial and egocentric learning and memory in rodent basic research and regulatory studies”. eng. In: Neurotoxicology and Teratology 45, pp. 75–90. issn: 1872-9738. doi: 10.1016/j.ntt.2014.07.003.

Wijnen, Kjell et al. (May 2024). “Rodent maze studies: from following simple rules to complex map learning”. eng. In: Brain Structure & Function 229.4, pp. 823–841. issn: 1863-2661. doi: 10.1007/s00429-024-02771-x.

Zou, Shimin and Chengyu Tony Li (Apr. 2019). “High-Throughput Automatic Training System for Spatial Working Memory in Free-Moving Mice”. In: Neuroscience Bulletin 35.3, pp. 389–400. issn: 1673-7067. doi: 10.1007/s12264-019-00370-z. url: https://pmc.ncbi.nlm.nih.gov/articles/PMC6527628/ (visited on 11/01/2025).

